# FLIPs: Novel Genetically Encoded Biosensors for Functional Imaging of Cell Signaling by Polarization Microscopy

**DOI:** 10.1101/2024.02.23.581811

**Authors:** Paul Miclea, Vendula Nagy-Marková, Robin Van den Eynde, Wim Vandenberg, Alina Sakhi, Alexey Bondar, Jitka Myšková, Peter Dedecker, Josef Lazar

## Abstract

Genetically encoded fluorescent biosensors convert specific biomolecular events into optically detectable signals. By revealing biochemical processes *in situ*, they have revolutionized cell biology. However, imaging molecular processes often requires modifying the proteins involved, and many molecular processes are still to be imaged. Here we present a novel, widely applicable design of genetically encoded biosensors that notably expand the observation possibilities, by taking advantage of a hitherto overlooked detection principle: directionality of optical properties of fluorescent proteins. The probes, which we term FLIPs, offer an extremely simple design, high sensitivity, multiplexing capability, ratiometric readout and resilience to bleaching artifacts, without requiring any modifications to the probe targets. We demonstrate their performance on real-time single-cell imaging of activation of G protein-coupled receptors (GPCRs), G proteins, arrestins, small GTPases, as well as receptor tyrosine kinases, even at endogenous expression levels. We also identify a new, pronounced, endocytosis-associated conformational change in a GPCR–β-arrestin complex. By demonstrating a novel detection principle and allowing many more cellular processes to be visualized, FLIPs are likely to inspire numerous future developments and insights.

## Main

Genetically encoded fluorescent biosensors allow visualizing specific cellular processes in living cells, tissues and animals. Such probes are often constructed by fusing a sensing protein to a fluorescent moiety (such as a fluorescent protein (FP)), creating a causal connection between the molecular process of interest and the fluorescent output[1, 2]. Usually, a conformational or structural change in the sensing protein is used to modulate one of several modes of fluorescence quenching, most often through facilitating (or restricting) access of water molecules to the fluorophore (e.g., in circularly permuted FPs), or through providing (or removing) a suitable excitation energy donor or acceptor (e.g., in bioluminiscence or fluorescence resonant energy transfer; BRET and FRET, respectively). The requirement for a sufficiently large conformational change in the sensing protein severely limits the number of phenomena that can be probed. Furthermore, many such biosensors require an insertion of a fluorescent moiety into the protein of interest (e.g. a receptor protein), leading to perturbations in its function. Although so-called translocation sensors, responding by redistribution of a fluorophore across the cell[3, 4], avoid the need for a conformationally-switching protein, their responses are hard to quantify and the number of their targets is limited.

Thus, there remains a strong need for new biosensors that would allow real-time *in vivo* observations of functional activity of a wide range of non-modified, endogenously expressed proteins, providing quantifiable, ratiometric readout while allowing multiplexing. In order to try to meet this need, we decided to exploit the well documented, but seldom utilized effects of optical directionality of fluorescent proteins.

Fluorescent molecules (including fluorescent proteins) behave like dipole antennas, with pronounced directional properties[5, 6]. Molecular directional properties can cause differences in rates of absorption of light of distinct linear polarizations (linear dichroism, LD), as well as fluorescence polarization. The two phenomena can report on molecular orientation (or orientational distribution) at the time of light absorption and emission by the fluorophore, yielding quantitative insights into protein structure and its dynamics not accessible by other means. Although observations of these phenomena offer intrinsically ratiometric readout, allow multiplexing (due to the need for only a single spectral band) and are relatively simple to implement, they have so far found only niche applications in biological microscopy[7–13]. This is because the existing biosensors based on optical directionality are of limited use: they modify the proteins of interest[14], require a specific design for each application, the observed responses are usually small[15], and efforts at improvements have largely been non-productive[8, 15]. For optical directionality to become a mainstream tool of biomolecular microscopy, these drawbacks would have to be overcome.

To address these challenges, we have conceived a novel class of molecular probes that we term FLIPs (**<u>F</u>**luorescence anisotropy and **<u>LI</u>**near dichroism **<u>P</u>**robes; see Acknowledgments) (Fig. 1a). A FLIP includes four distinct functional components: a fluorescent moiety (i) (such as an FP), bearing at one of its termini a flexible linker (ii) and a membrane-targeting moiety (iii), and at the other terminus a moiety (iv) capable of binding the target of the probe. We hypothesized that in absence of the probe’s target, the FP moiety would adopt a largely random molecular orientation, while in presence of the target, the orientation of the FP with respect to the cell membrane would become restricted.

**Figure 1:**
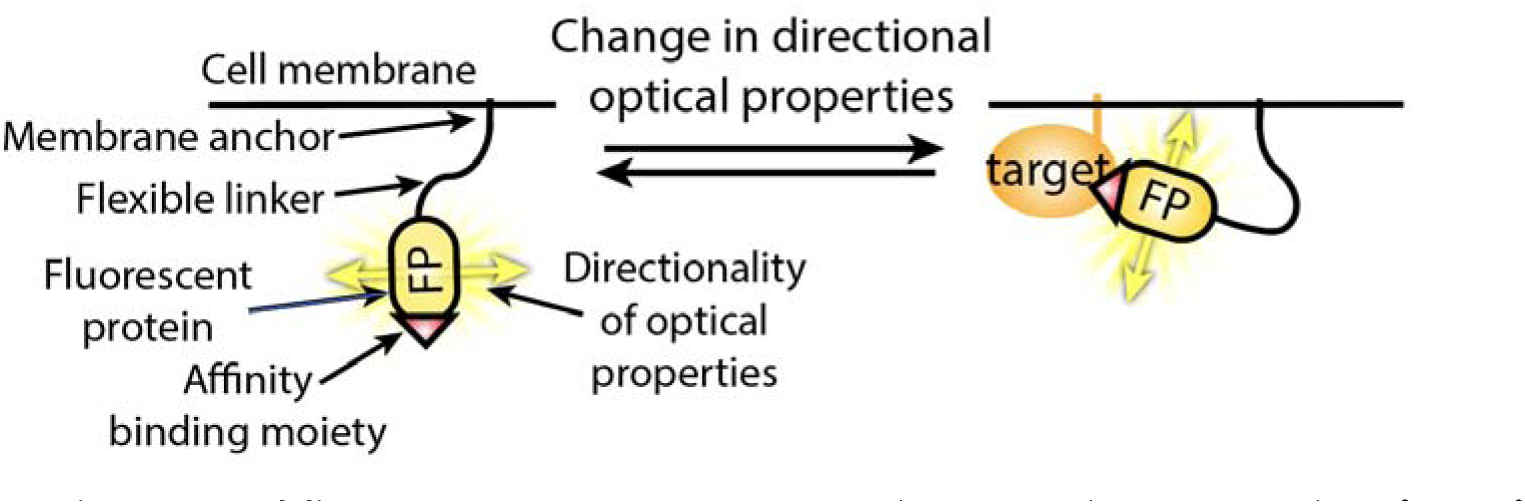
General design of fluorescence anisotropy and linear dichroism probes (FLIPs).

The FLIP design seems to address all of the above-mentioned drawbacks of existing biosensors based on optical directionality. The target molecules do not need to be modified in any way. By virtue of its clear molecular mechanism of function, simplicity, modularity and versatility, the FLIP architecture promises to allow rapid generation and optimization of a broad range of sensitive biosensors. The expected robust, predictable changes in molecular orientation of the fluorescent moiety should yield strong observable signatures, while retaining inherent advantages (insights into protein structure and dynamics, ratiometric output using a single spectral channel, resistance to bleaching artifacts). Here we present the results of our investigations of the proposed FLIP biosensors.

### Initial FLIP designs

For our initial attempts (Fig. 1), we chose to develop a FLIP that would report on the presence of the activated form of the G protein Gαi1. Since Gαi1 signaling plays an important role in physiology (transducing signals from numerous GPCRs), it had previously been investigated both by polarization microscopy[8, 14, 15] and other methods[2, 16], allowing comparisons. As the affinity binding moiety we chose the peptide KB1753, known to bind activated Gαi subunits[16, 17]. The enhanced green fluorescent protein (eGFP) served as the fluorescent moiety. We used the hexapeptide Gly-Ser-Gly-Gly-Ser-Gly as the flexible linker. Finally, as membrane anchors[18] we tested the N-terminal palmitoylation signal peptide derived from GAP43, the transmembrane domain of the interleukin receptor IL4Rα, the C-terminal palmitoylation/farnesylation domain of H-Ras, and the C-terminal farnesylation domain of K-Ras.

To determine whether the four probes (GAP43-eGFP-KB1753, IL4R-eGFP-KB1753, KB1753-eGFP-HRas, KB1753-eGFP-KRas) interact with the intended target, we observed their linear dichroism in living HEK293 cells, in absence and in presence of a constitutively active mutant of Gαi1 (denoted as Gαi1*). The results are illustrated in Fig 2, Suppl. Figs 1-3, and summarized in Table 1. With the exception of GAP43-eGFP-KB1753, we observed striking differences in LD between cells expressing only a probe and those co-expressing the probe and Gαi1*. The effect was most pronounced in IL4Rα-eGFP-KB1753 which, however, showed irregular membrane localization (Suppl. Fig. 2). Thus, we selected KB1753-eGFP-HRas for further development and testing. Modifying the length of the flexible linker brought little benefit, as probes with longer linkers exhibited smaller changes in LD, and, ultimately, poorer cell membrane localization (Suppl. Fig. 3).

**Figure 2:**
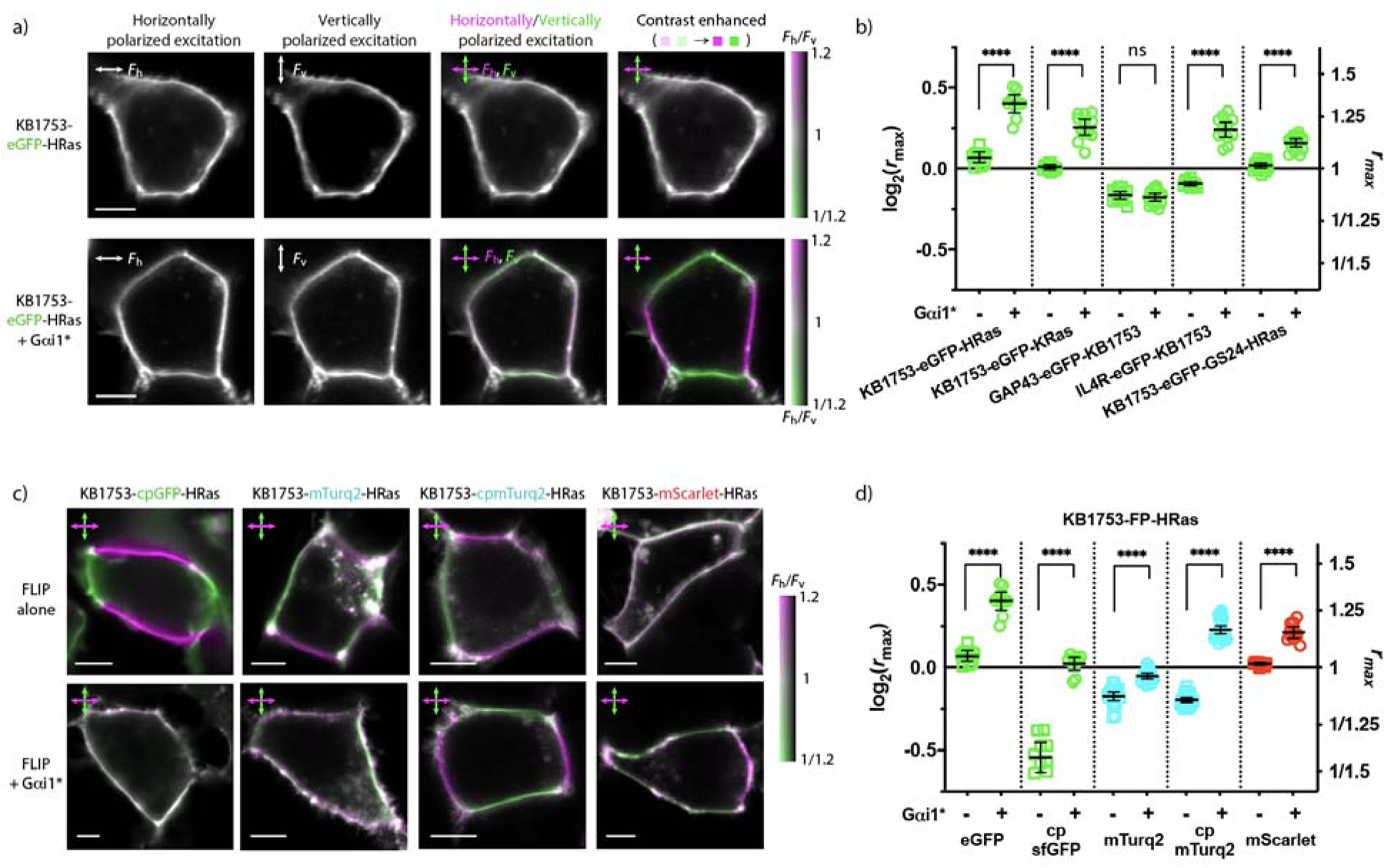
Detecting the presence of a constitutively active mutant of Gαi1 (Gαi1*) by FLIP and LD microscopy. **a)** A HEK293 cell expressing KB1753-eGFP-HRas (top) and coexpressing KB1753-eGFP-HRas and Gαi1* (bottom). From left to right: image acquired using excitation light polarized horizontally; image acquired using excitation light polarized vertically; the two preceding images colored magenta and green, respectively, and overlaid; same as in the preceding image, but the coloring adjusted to cover the shown range of dichroic ratios (F_h_/F_v_,), as indicated by the color bar. In cells expressing Gαi1*, the FLIP exhibits pronounced LD. Scale bars: 5 µm. **b)** LD of eGFP-bearing FLIPs containing various membrane anchors and linkers, in absence and presence of Gαi1*. Means and 95% confidence intervals are indicated. With the exception of GAP43-eGFP-KB1753, all tested FLIPs exhibit significantly different LD in presence of Gαi1*. **c)** Representative LD microscopy images of HEK293 cells expressing FLIPs of the KB1753-FP-HRas design, bearing various FPs. Coloring as in **a**. Scale bars: 5 µm. Top: cell expressing the indicated FLIP; bottom: cells co-expressing the indicated FLIP and Gαi1*. Differences in LD between cells expressing only a FLIP and co-expressing the FLIP and Gαi1* are apparent. Notably, cells expressing KB1753-cpmTurq2-HRas exhibit LD of a different sign in absence and presence of Gαi1* (inversion of the magenta-green pattern). **d)** LD of FLIPs of the KB1753-FP-HRas design, bearing different FP moieties. Means and 95% confidence intervals are indicated. Presence of Gαi1* had a significant effect on the LD of all tested FLIPs. Marked differences between the distinct FLIPs are apparent.

**Figure 3:**
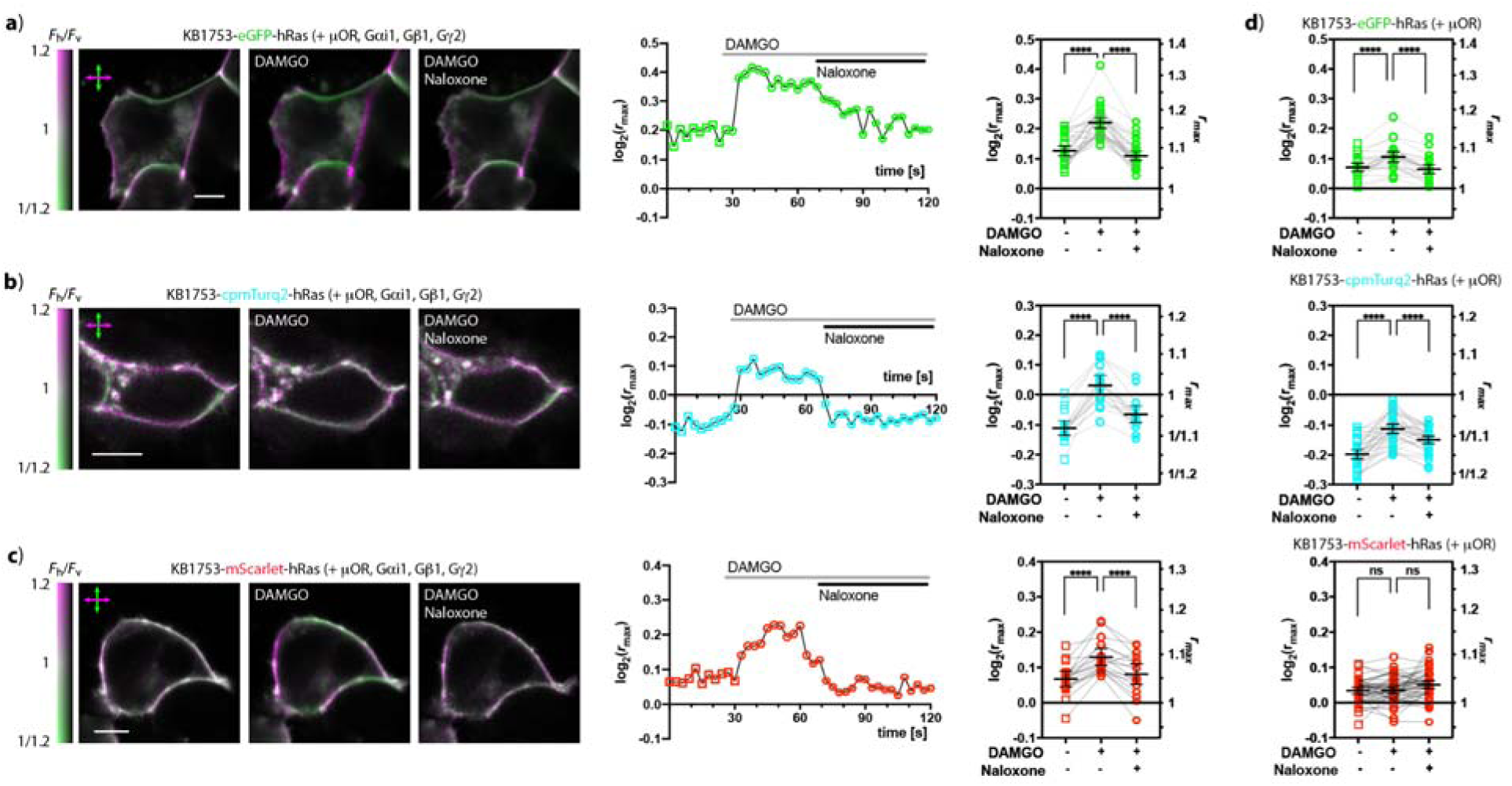
Imaging activity of Gαi1 by LD microscopy and FLIPs of the KB1753-FP-HRas design. **a-c)** HEK293 cells overexpressing the µ-opioid receptor (µOR), Gαi1, Gβ1, Gγ2 and the indicated FLIP, imaged before and after addition of a µOR agonist (DAMGO), and after addition of an excess µOR antagonist (naloxone). **a)** KB1753-eGFP-HRas; **b)** KB1753-cpmTurq2-HRas; **c)** KB1753-mScarlet-HRas. Left: LD microscopy images showing a cell before addition of DAMGO, after addition of DAMGO, and after addition of naloxone. All scale bars 5 µm. Middle: temporal profile of LD in response to µOR activation and inactivation of the shown cell. Right: LD of cell prior to activation, 30 s after addition of DAMGO, and 30 s after addition of naloxone. Change in LD in response to addition of DAMGO and naloxone are apparent for all three FLIPs. **d)** LD of HEK293 cells overexpressing µOR and the indicated KB1753-FP-HRas (not overexpressing any G protein), before and after addition of DAMGO and after addition of naloxone. Mean values and 95% confidence intervals indicated.

Replacing the eGFP moiety with other FPs (Fig 2c, d) affected the directional optical properties of the KB1753-FP-HRas constructs, sometimes in unexpected ways. Predictably, modifying the orientation of the fluorophore with respect to the cell membrane by circular permutation of the FP moiety (cpsfGFP[19], cpmTurq2[20]), affected the LD of the respective sensors, both in absence and presence of Gαi1*. To our surprise, some FLIPs (namely KB1753-cpsfGFP-HRas and KB1753-mTurq2-HRas) showed higher LD in absence of Gαi1* than in its presence. Some highly similar constructs (KB1753-eGFP-HRas and KB1753-mTurq2-HRas) exhibited opposite Gαi1*-induced behaviors. Strikingly, KB1753-cpmTurq2-HRas showed pronounced LD both in absence and in presence of Gαi1* - but with opposite signs. Without detailed investigations, we attribute the observed idiosyncratic behaviors of these constructs to sequence-specific interactions between the FP moiety and the plasma membrane. In the future, such interactions may be exploited in rational development of improved FLIPs.

Importantly, our results demonstrate that FLIPs can accommodate a wide range of FPs spanning the whole visible light spectrum. All tested FPs were capable of generating a strong LD contrast, albeit sometimes in unexpected fashion. A particularly notable kind of contrast was observed in KB1753-cpmTurq2-HRas, which showed opposite signs of LD in absence and in presence of Gαi1*, allowing simple, unambiguous visual identification of different conformational states of the FLIP even without LD quantitation.

### Observing activation of Gαi1

We then explored whether the FLIPs that showed distinct LD in absence and presence of Gαi1* can also report on activation and inactivation of non-modified Gαi1 (Fig. 3). To ensure close to equimolar concentrations, we overexpressed a KB1753-based FLIP, as well as non-modified G proteins Gαi1, Gβ1, Gγ2 and the µ-opioid receptor (µOR). Prior to activation of the receptor, all three tested FLIPs (KB1753-eGFP-HRas, KB1753-cpmTurq2-HRas and KB1753-mScarlet-HRas) exhibited LD somewhat higher than that of the probes overexpressed alone (Fig. 3). This is likely due to basal activity of µOR and other Gi-coupled GPCRs present in the cells, as well as non-specific binding of KB1753 to the non-activated form of Gαi1. Upon addition of DAMGO, a full µOR agonist[21], the LD of all three probes quickly changed, in a direction consistent with our observations of the effects of Gαi1*. Cells expressing KB1753-cpmTurq2-HRas often changed the sign of LD (LD inversion, or LD ‘flip’), allowing easy visual identification of different functional states of Gαi1. The responses of all three FLIPs were smaller than what could be expected from our experiments with Gαi1*, likely due to incomplete activation of the Gαi1 pool by µOR. Upon addition of an excess of a µOR antagonist (naloxone)[16], the LD of all three FLIPs returned to values close to those observed prior to µOR activation, with residual responses attributable to continuous presence of DAMGO. The observed responses to µOR activation/inactivation suggest that the binding of the FLIP to the activated Gαi1 does not substantially affect the kinetics of the G protein activation or inactivation. Our results show that FLIPs can be used for real-time single-cell imaging, with subcellular resolution, of activation and inactivation of non-modified, overexpressed G proteins, exceeding the abilities of other sensors.

To investigate whether FLIPs can report on activity of endogenously expressed Gαi proteins, we observed cells overexpressing a FLIP and µOR (allowing reliable Gαi activation) but expressing G proteins endogenously (Fig. 3d). Our results show that the LD of KB1753-eGFP-HRas and KB1753-cpmTurq2-HRas changes significantly both upon µOR activation by DAMGO and inactivation by naloxone. Unlike the other two FLIPs, KB1753-mScarlet-HRas showed no statistically significant changes in LD upon activation/inactivation of endogenously expressed Gαi, even after observing over 50 cells.

The LD changes observed in cells overexpressing µOR and KB1753-eGFP-HRas or KB1753-cpmTurq2-HRas are in line with those we observed in cells that also overexpressed Gαi1, Gβ1 and Gγ2, but markedly smaller. Based on our measurements of LD in cells expressing Gαi1*, we estimate that about 15 % of the FLIP molecules become bound to the endogenous Gαi upon activation. In theory, reducing the molar excess of the FLIP over the endogenous Gi might lead to a larger fraction of the FLIP becoming bound upon Gi activation, and in turn, to a larger and more easily discernible change in LD. However, since we did not notice any relationship between the magnitude of the LD change and the FLIP expression level, we believe that the potential benefit of a lower FLIP concentration is being negated by the relatively low affinity of KB1753 for the activated Gαi subunits (Kd = ∼1 µM)[17]. Improving the sensitivity of detection of endogenous G protein signaling will likely require increasing the affinity of the FLIP binding moiety. Despite the limits imposed by the affinity of KB1753, our ability to image activation of non-modified, endogenously expressed G proteins exceeds the best existing approaches[2], and is likely to improve further with future developments.

### Observing activation of GPCRs

Encouraged by our success in observing activity of Gαi1, we explored expanding the range of FLIP applications to observations of activity of GPCRs. The versatility of the FLIP design allowed us to create biosensors based on nanobodies NB33 and NB80, which specifically bind the activated forms of µOR and the β2-adrenergic receptor (β2AR), respectively[3, 22] (Fig. 4). Briefly, the LD of NB33-eGFP-HRas exhibited rapid LD inversion upon addition of DAMGO. The response could be reversed by application of naloxone. NB80-eGFP-HRas responded to activation of β2AR (by isoproterenol) by rapid LD inversion. In fact, the observed LD change was so fast (< 100 ms) and pronounced that it could often be observed during the acquisition of a single frame by the laser scanning microscope (Fig. 4c). In contrast, the observed rate of β2AR inactivation by a β2AR inverse agonist (propranolol) was on the order of minutes. We reasoned that the high binding affinity of NB80 for β2AR (reported to be 3 nM for the isoproterenol-activated receptor[3]) was preventing the receptor from leaving the active conformation. Thus, guided by the known structure of the β2AR-NB80 complex[23] (PDB ID 3P0G), we mutated four amino acids of NB80 that form contacts with the active form of β2AR, but away from the loop that fits into the central cavity of the activated receptor (Fig 4d). The resulting FLIP (NB80Lite-eGFP-HRas) exhibits optical properties similar to NB80-eGFP-HRas, but virtually no affinity for the non-activated β2AR, and a much faster response to β2AR inactivation (a few seconds). The large magnitude of the probe’s responses preserves the ability to unambiguously recognize the state of the β2AR by imaging over a period of tens milliseconds (as shown in Fig 4c for NB80-eGFP-HRas) or, if scanning a small area, over tens of microseconds.

**Figure 4:**
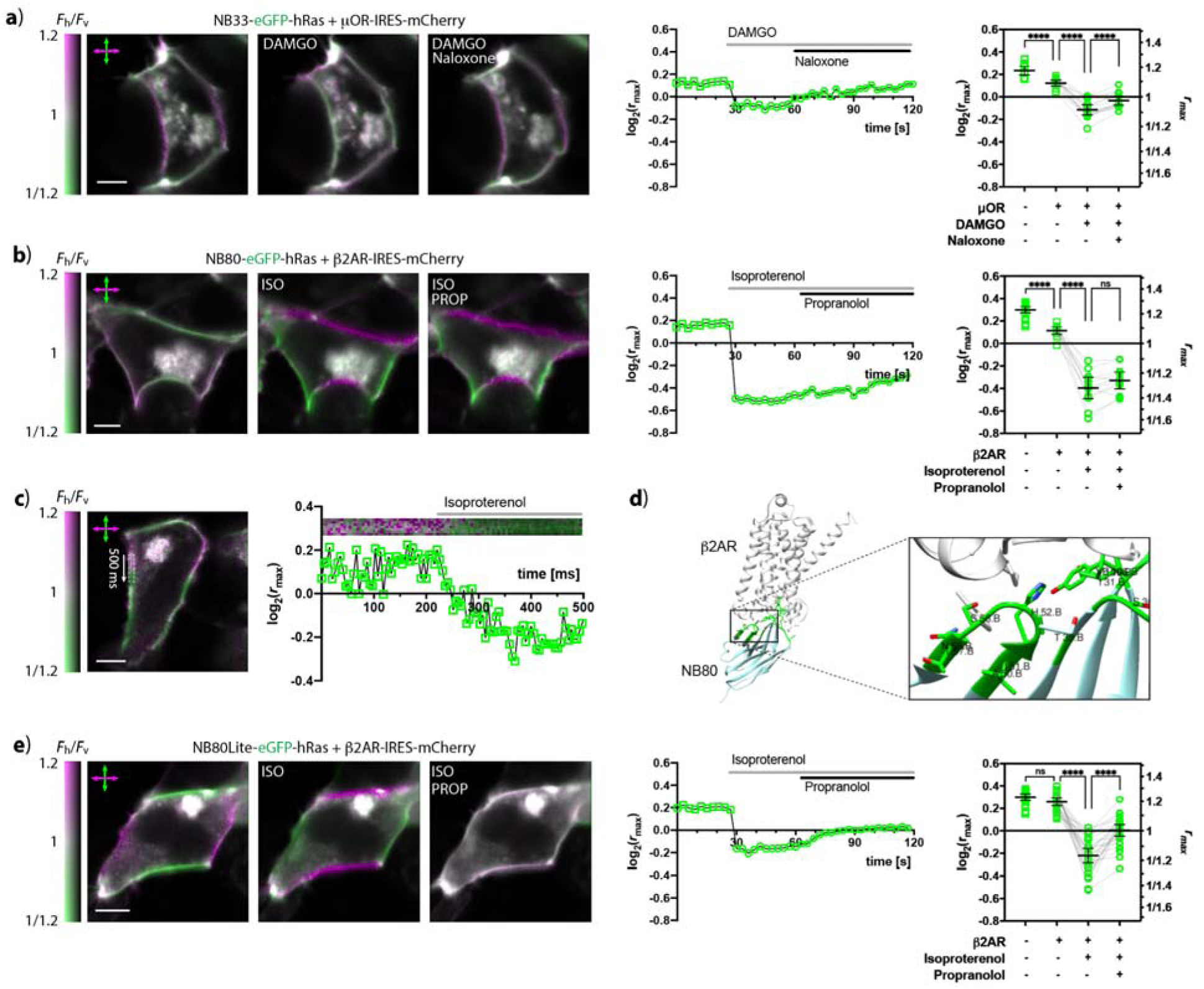
Imaging GPCR activity by LD microscopy and FLIPs. **a)** Imaging µOR activity in HEK293 cells overexpressing µOR-IRES-mCherry and NB32-eGFP-HRas. Left: A cell observed before stimulation, after µOR activation (by addition of DAMGO), and after µOR inactivation (by addition of naloxone); Middle: temporal profile of LD during µOR activation and inactivation of the shown cell; Right: values of LD observed in absence of µOR, and in its presence before stimulation, 30 s after activation, and 30 s after inactivation (each data point represents a single cell; mean values and 95 % confidence intervals indicated). **b)** β2AR activity in HEK293 cell overexpressing β2AR-IRES-mCherry and NB80-eGFP-HRas; isoproterenol (ISO) and propranolol (PROP) were used for β2AR activation and inactivation, respectively. **c)** An NB80-eGFP-HRas response during acquisition of a single image. Left: an image of a cell imaged during agonist delivery, showing an LD inversion occurring while the image was being acquired; Right: a temporal profile of the LD inversion in the indicated region of the image on the left. **d)** Structure of the β2AR-NB80 complex (PDB ID 3P0G) showing the loop region of NB80 mutated to create NB80Lite. Residues of NB80 forming contacts with β2AR are shown in green. **e)** Imaging β2AR activity using NB80Lite-eGFP-HRas. All scale bars 5 µm.

Our results demonstrate that FLIPs can be used for sensitive, real-time observations of activity of non-modified GPCRs. They also show that nanobodies can serve as affinity moieties in FLIPs, while highlighting the importance of a suitable binding affinity of the FLIP for its target. Lowering the binding affinity by mutating key residues yielded a probe capable of fast and sensitive detection of both β2AR activation and inactivation. In combination with our observations of KB1753-based probes, our results with NB80-based FLIPs place limits on the range of binding affinities suitable for FLIP applications (likely 10-100 nM, depending on the target concentration).

### Observing activity of other proteins

To explore the versatility of the FLIP design and a range of applications, we developed FLIPs containing the RH domain of GRK2 (known to bind the activated form of Gαq[24]), nanobody NB35 (known to bind the activated form of Gαs[23]); the C-terminal domain of GRK2 (known to bind the Gβγ dimer often released upon activation of G protein heterotrimers[24]), PAK1 (known to bind the activated form of the small GTPase RacA[25]), as well as a recently developed GP2-derived small affinity protein known[26] to bind to the ectodomain of the insulin receptor 1 (IR1) (Fig. 5). Presence of their respective targets generally increased LD (that is, it increased the absolute values of the observed dichroic logratios), consistent with lowering the orientational freedom of the fluorescent moiety. Interestingly, GP2(1)-meGFP-CD59, a probe of activation of the insulin receptor, showed intracellular localization in absence of its target, despite containing a leader sequence and a GPI anchor (derived from the protein CD59[27]). In contrast, the probe localized to the cell membrane in cells overexpressing both the probe and IR1. Upon addition of insulin, the LD of the probe decreased, consistent with insulin-induced conformational changes in the IR1 ectodomain.

**Figure 5:**
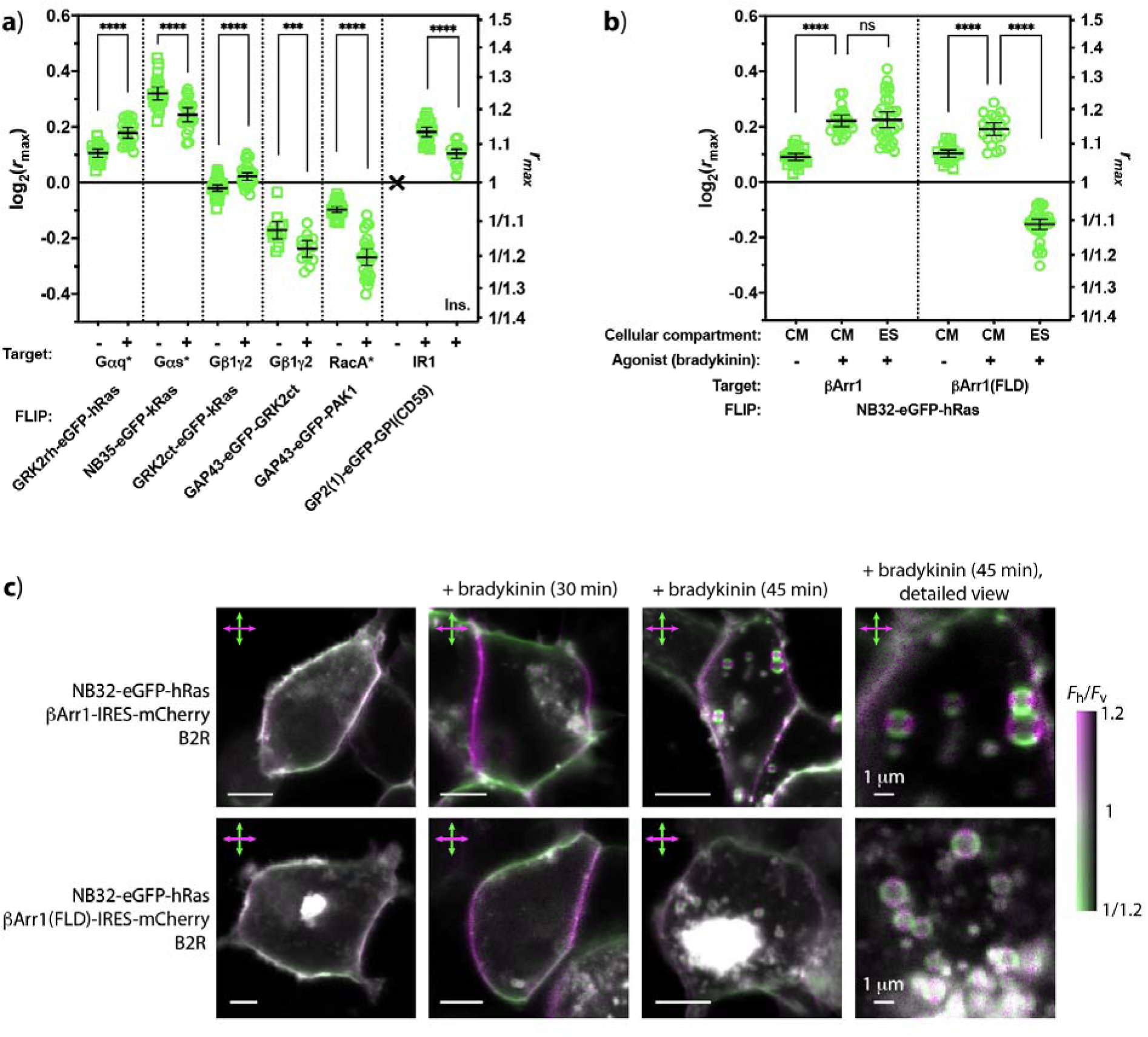
Other FLIPs. **a)** Results of LD microscopy observations of various FLIPs, in absence and in presence of their targets, namely the Gα subunits Gαq and Gαs, the Gβ1γ2 dimer, the small GTPase RacA, and the insulin receptor 1 (IR1). Constitutively active mutants indicated by an asterisk (*). Stimulation by insulin (Ins.) indicated. Each data point represents a single cell; a cross (x) indicates poor membrane localization; mean values and 95 % confidence intervals are shown. **b)** Observations of NB32-eGFP-hRas in response to arrestin activation/recruitment due to stimulation of the bradykinin receptor (B2R) by bradykinin. Cells overexpressing β-Arrestin1 (βArr1) and its finger-loop deletion mutant βArr1(FLD) exhibit similar LD responses in the cell membrane (CM), but markedly different responses in endosomes (ES) that emerge upon prolonged activation. **c)** Images of typical cells overexpressing B2R, NB32-eGFP-hRas, and βArr1 (top row) or βArr1(FLD) (bottom row). From left to right: before application of bradykinin; 30 min after application of bradykinin; 45 min after application of bradykinin; and a detailed view showing endosomes, 45 min after application of bradykinin. Scale bars: 5 µm unless indicated otherwise.

These observations illustrate the potential range of FLIP applications, which include studies of various G protein activities (Gαq, Gαs, Gβγ), function of small GTPases (activation of RacA, critical for embryonic development), or activity of receptor tyrosine kinases (i.e., IR1, of crucial importance in diabetes). The results also show how wide a range of molecules can serve as binding moieties in FLIPs: peptides, nanobodies and nanobody-like affinity scaffolds, but also natural binding partners (GRK2, PAK1) of the target proteins. Conveniently, by virtue of being natural binding partners of endogenously expressed targets, such molecules are likely to exhibit affinities suitable for observations of targets expressed endogenously. Thus, our results open an avenue to observations of an exceedingly wide range of non-modified, endogenously expressed signaling molecules.

### Endocytosis-associated rearrangement of GPCR–arrestin complexes

Among valuable potential FLIP targets are also arrestins. Arrestins play an important role in GPCR signal transduction, complementing or competing with signaling by G proteins[28, 29]. The roles of G proteins and arrestins are compartmentalized, both spatially (cell membrane vs. endosomes) and temporally (so-called first vs. second wave of GPCR signaling)[30]. Upon GPCR activation, arrestins can bind to the receptor’s phosphorylated C-terminus (partial engagement), or to both the C-terminus and the GPCR’s central cavity (full engagement)[31, 32]. Partial arrestin engagement might allow formation of GPCR–G protein–arrestin supercomplexes, whose existence has, however, only been demonstrated with purified, reconstituted proteins, under highly favorable conditions[33]. In contrast, our approach promised real-time observations of arrestin molecular interactions in living cells, with subcellular resolution.

In order to detect interactions of arrestins with GPCRs, we prepared NB32-eGFP-HRas[34]. This FLIP utilizes NB32, a nanobody specific for activated arrestins bound to the phosphorylated C-terminus of a GPCR[31]. To stimulate arrestin recruitment, we used the bradykinin receptor 2 (B2R) [4, 35, 36]. Cells overexpressing non-modified B2R, β-arrestin 1 (βArr1) and NB32-eGFP-HRas responded to activation by bradykinin by an increase in LD of the probe in the cell membrane (Fig. 5b, c), likely corresponding to the formation of the fully engaged B2R–βArr1 complex. Membrane vesicles (endosomes), which formed upon internalization of the activated B2R, exhibited LD similar to that observed in the cell membrane of the activated cells. Thus, NB32-eGFP-HRas can detect activation of βArr1 both in the cell membrane and in endosomes.

To test whether NB32-meGFP-hRas can distinguish between different modes of arrestin binding to activated GPCRs, we prepared a βArr1 mutant (βArr1(FLD)), bearing a deletion of the finger-loop region. Such deletions have been shown[31] to augment arrestin’s partial engagement with GPCRs, reducing the formation of fully engaged GPCR–arrestin complexes. In presence of βArr1(FLD), activation of B2R leads to an increase in LD of NB32-eGFP-HRas present in the cell membrane similar to that observed in presence of non-mutated βArr1. However, strikingly, upon internalization into endosomes, the LD of the probe undergoes an inversion. This is in contrast to the behavior observed with non-mutated βArr1, and likely corresponds to formation of partially-engaged complexes of B2R with βArr1(FLD).

The endocytosis-associated LD inversion observed with βArr1(FLD) is robust and highly reproducible, and seems to occur irrespective of the endosome size (for vesicles of diameters 0.5 – 2 µm; Suppl. Fig. 4), cellular localization (periplasmic/intracellular; Fig. 5c), expression levels (weakly/brightly fluorescent; Fig. 5c) or vesicle age (nascent/mature). Possible causes for the observed conformational change include B2R’s interactions with the caveolin-[37, 38] or clathrin-associated[39, 40] endocytosis machinery, effects of the endosomal lipid composition[41], pH, membrane curvature[42], or a yet-to-be-identified process occurring during the formation of endosomes. Although all of these mechanisms are expected to act both on B2R–βArr1 and B2R–βArr1(FLD), we could only observe rearrangement in the latter. This might reflect a true difference in behaviors between βArr1 and βArr1(FLD). Our observations suggest that a rearrangement occurs in complexes between GPCRs and βArr1 of suitable affinities[4], opening a possibility of downstream signaling bifurcation between distinct GPCRs. Intriguingly, the endosomal localization of the observed conformational transition is consistent with proposed mechanisms[33] of second-wave GPCR signaling that involve supercomplexes GPCR–G protein–arrestin.

Our observations of a previously unseen endocytosis-associated rearrangement of a GPCR–arrestin complex demonstrate the importance of capacity of FLIPs and LD microscopy to visualize interactions between non-modified GPCRs and arrestins with subcellular resolution. Importantly, they also illustrate a unique ability to distinguish between distinct forms of protein-protein interactions. This ability, combined with the fact that the FLIP targets bear no modifications, opens doors to revealing hidden aspects of cell signaling, under conditions markedly closer to natural than previously possible.

## Conclusions

Our results demonstrate the potential of using optical directionality of fluorescent molecules for design of genetically encoded fluorescent biosensors. Despite being known for decades[5], optical directionality of fluorescent proteins has remained largely overlooked as a useful detection principle, due to lack of suitable molecular designs. By identifying FLIPs, a large, widely applicable and exceedingly simple class of molecules capable of reporting on numerous scientifically and pharmacologically important molecular processes of cell signaling, we hope to open a new avenue to developing better genetically encoded biosensors.

The FLIP design offers a powerful combination of desirable properties. The molecular targets do not need to be modified in any way, and the approach is sensitive enough to detect even activity of endogenously expressed proteins. A single FLIP can report on the presence or absence of a target, but, uniquely, it can also distinguish among several states of the target, yielding more information than other types of biosensors. This ability allowed us to unambiguously identify a pronounced conformational change in GPCR-arrestin complexes and to show that it occurs specifically in endosomes. Our results bring a new insight into mechanisms of signaling by GPCRs, an exceedingly large and important class of proteins. The ability to perform such observations in living cells, in real time, in signaling proteins that have not been modified in any way, sets a new standard in microscopy imaging of protein signaling.

The FLIP design does present some limitations. LD observations rely on the shape of the cell membrane, which can be irregular. However, due to the mathematical relationship between LD and cell membrane orientation, accurate log_2_(*r*_max_) determinations can be made[43] even using a very small suitable section of a cell membrane. Another apparent FLIP limitation is the requirement for the target to be membrane-associated. However, suitably designed FLIPs should respond even to presence or absence of a cytoplasmic target, rendering this reservation largely mute. It could be argued that FLIPs are partly interchangeable with translocation-based biosensors. However, protein translocation is slower, difficult to accurately quantify, does not yield structural information about the biosensor, and, therefore, a single translocation biosensor cannot distinguish among multiple states of the target. In fact, translocation biosensor uses are more strictly limited to membrane-associated targets than those of FLIPs. Finally, by binding to their targets, FLIPs might sequester the targets from downstream signaling. Nonetheless, choosing a suitable combination of FLIP concentration and binding affinity should allow overcoming this drawback. Therefore, FLIPs, in combination with polarization microscopy, present a substantial, widely applicable improvement over existing approaches.

In order to observe FLIPs, various polarization microscopy implementations can be used. Our preferred FLIP imaging modality is linear dichroism microscopy, implemented on a laser-scanning confocal microscope[43]. This implementation combines the advantages of optical sectioning, a wide range of detection channels, and relatively (in comparison with fluorescence polarization microscopy) low depolarization by the objective lens and by the excited FP molecules[44]. Nonetheless, our results show that FLIPs can also be used with fluorescence polarization microscopy. Combining FLIPs with various other polarization microscopy modalities (including wide-field[45], multipoint[46], holographic[47], or super-resolution[48]), as well as with a growing range of publicly available software tools for polarization microscopy image analysis[43, 49–52], promises to allow functional imaging in an excitingly wide range of biological contexts, from single-molecule studies in individual organelles or cellular compartments to large scale observations in intact tissues or animals.

By allowing innumerable combinations of affinity binders, anchoring moieties, linkers and fluorescent molecules (potentially including self-labeling proteins), the FLIP design allows rapid development of a wide variety of new molecular biosensors. By drawing attention to optical directionality of fluorescent proteins, our results will likely inspire other biosensor designs that take advantage of this fundamental property of fluorescent molecules. We hope that the present work will ultimately transform polarization-resolved fluorescence microscopy, a simple, long overlooked optical microscopy modality, into a mainstream tool of functional biomolecular imaging.

## Methods

### Constructs

The plasmids encoding KB1753-eGFP-HRas and GAP43-eGFP-KB1753 (both in the pcDNA3.1(+) backbone) were prepared by gene synthesis (GenScript). The constructs were designed to allow excision of the DNA encoding the affinity binding moiety, the FP moiety, and the membrane-targeting moiety by a unique combination of restriction endonucleases (EcoRI/XhoI, XhoI/BamHI and BamHI/NotI, respectively). Other FLIPs were prepared from the two starting plasmids by a suitable restriction digestion, followed by ligation of the desired DNA fragment, obtained through gene synthesis (GenScript). In some cases, preliminary experiments were performed using constructs kindly provided by A. Shukla (plasmids containing NB32, βArr1, βArr1(FLD), B2R) and M. von Zastrow (NB33). As fluorescent proteins we used eGFP bearing the dimerization-preventing mutations A206K, L221K and F223R. Circularly permuted FPs (cpmTurq2, cpsfGFP) were based on superfolder mTurquoise2 and sfGFP2. FLIP targets were cloned into a pcDNA3.1(+) plasmid backbone containing either IRES-mCherry or IRES-mTurquoise2, allowing identification of target-expressing cells by fluorescence of the bicistronically expressed FP, while the targets themselves bore no modifications. Constitutively active mutants (Gαi1(Q204L)), Gαq(Q209L), Gαs(Q227L), RacA(Q61L)) were prepared by PCR mutagenesis. NB80Lite was created by introducing mutations H52S, S56G, T57S and N58A into NB80-eGFP-HRas by PCR mutagenesis.

### Mammalian cell culture

HEK293 cells obtained from ATCC, passaged less than 15 times were used throughout. Cells were regularly tested for mycoplasma by Hoechst staining. For imaging, cells were plated in 8- or 18-well imaging slides (iBidi) and transfected with the desired plasmid (0.5 µg per well) using Lipofectamine 3000 (Thermo Fisher) and the manufacturer’s procedure. Cells were observed by LD microscopy 1 or 2 days after transfection. During imaging, GPCR agonists (DAMGO, isoproterenol, bradykinin; final concentrations 10 µM) and antagonists (naloxone, proterenol; final concentrations 100 µM) were added by manual pipetting. Agonists and antagonists were purchased from Sigma-Aldrich.

### Microscopy

LD imaging was carried out as described previously[8, 43], on a customized laser scanning confocal microscope (Olympus FV1200), equipped with a polarization modulator (RPM-2P, Innovative Bioimaging, L.L.C.) alternating the excitation light linear polarization between acquisition of subsequent pixels by the microscope. The objective (UApoN340, NA1.15, Olympus, Japan) and pixel dwell times of 10 µs were used throughout. Cells expressing constructs bearing mTurqoise2 were imaged using 405 nm illumination and a 450-550 nm emission window. Cells expressing constructs containing eGFP were imaged using 488 nm illumination, and emission 510-610 nm was collected. Cells expressing constructs with mScarlet and mCherry were imaged using 543 nm excitation; emission 560-650 nm was collected. The resulting images were processed and analyzed as described[43], using Fiji and publicly available macros for LD image analysis, as well as Polaris+ software (Innovative Bioimaging, L.L.C.). Briefly, each raw image (containing information on fluorescence intensity acquired with both excitation polarizations) was deconvolved into a pair of images, each holding information on fluorescence intensity acquired with one polarization of the excitation light. For LD visualization, the images acquired with horizontal and vertical polarization of the excitation light were colored magenta and green, respectively, and the color saturation was adjusted to match the desired range of LD. To quantitate LD, parts of images containing the cell membrane were semi-automatically segmented, and the shape of the cell outline was approximated by a spline. For each segmented pixel, the dichroic logratio (log_2_(*r*) = log_2_(*F*_h_/*F*_v_)) and membrane orientation (angle θ; slope at the nearest point on the spline approximating the cell outline) were determined as described previously[43]. At least ten cells were observed and analyzed for each construct or combination of constructs, over at least three different weeks. Distributions of the log_2_(*r*_max_) values for a large majority of our experiments were normal (according to Anderson-Darling, D’Agostino & Pearson tests). To maintain consistency, we treated all log_2_(*r*_max_) values as normally distributed.

Fluorescence polarization imaging was carried out using a modified Olympus FV1200 confocal microscope. For illumination, the optical fiber with a collimator was detached from the microscope body, and a Glan-Taylor polarizer (GT10-A, Thorlabs) and an achromatic quarter-wave plate (WP140HE, Edmund Optics) oriented to ensure circularly polarized sample illumination were placed in the laser beam path before the beam was steered into the scanning unit. In the detector unit, a dichroic beam splitter assembly was replaced by a polarizing beam splitter cube (PBS201, Thorlabs) flanked by sheet polarizers (XP42, Edmund Optics, extinction ratio 9000:1). To allow comparisons with observations of linear dichroism, the extent of fluorescence polarization was quantified analogously, by observing the logratio of the two fluorescence intensities along the cell outline and determining, by fitting, the maximum value (log_2_(*r*_max_)), corresponding to the value of log_2_(*F*_||_/*F*_⊥_) at membrane sections oriented horizontally in the image (θ = 0).

## Acknowledgements

The research was supported by Ministry of Education, Youth and Science of the Czech Republic grant LTAIN19167 (J.L.) and EIC PathFinder project UniSens (J.L.). Data storage was provided by the CESNET LM2015042 facility under the program “Large Research, Development, and Innovation Infrastructures.” We appreciate the help of Z. Vaitova with preparing constructs and performing microscopy imaging and the help of P. Khoroshyy with microscopy equipment. We are grateful to P. Dedecker (KU Leuven) for support and for editing the manuscript. We thank A. Shukla (IIT Kanpur) and members of his laboratory for providing constructs and for insightful discussions. We thank M. Garcia-Marcos (U. Boston) for providing constructs for preliminary experiments. We appreciate that M. von Zastrow (UCSF) provided constructs that served in FLIP development. We thank M. Sommer (ISAR Bioscience) for reading the manuscript and providing helpful suggestions. The terms ‘fluorescence anisotropy and linear dichroism probes’, ‘FLIP’ and ‘FLIPs’ were contributed by Innovative Bioimaging, s.r.o., which may claim copyright.

## Author contributions

P. M. designed and prepared constructs, carried out microscopy imaging, data processing and analyses, and contributed to writing the paper. V.M. performed imaging, data processing and analyses. A.S. developed and investigated FLIPs of IR activity. A.B. inspired the idea of FLIPs and supervised experiments. J.M. performed optical microscopy and image analyses. J.L. formulated, directed and managed the project, provided technical expertise, designed constructs, and wrote the manuscript, with input from the other co-authors.

## Microscopy data summary

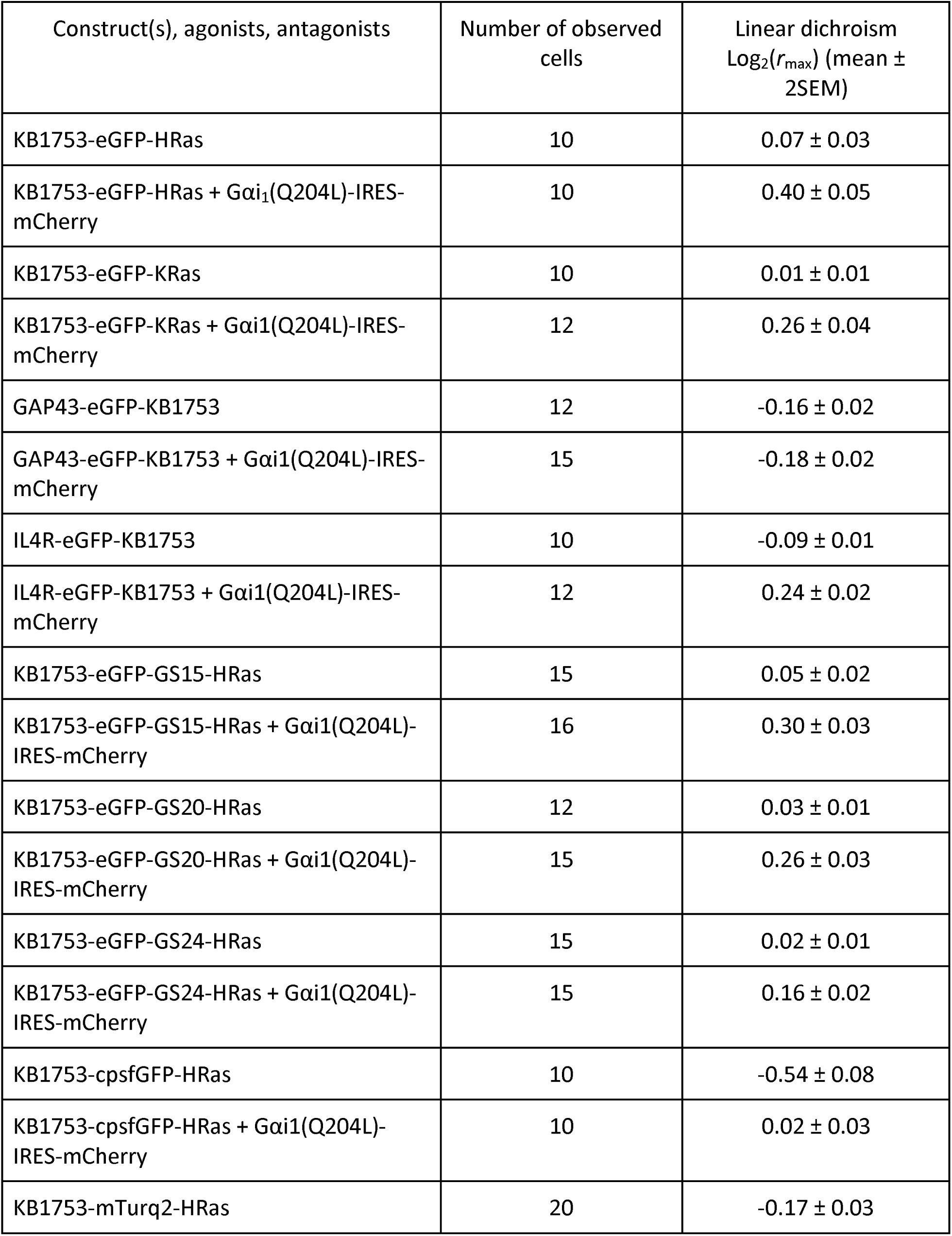
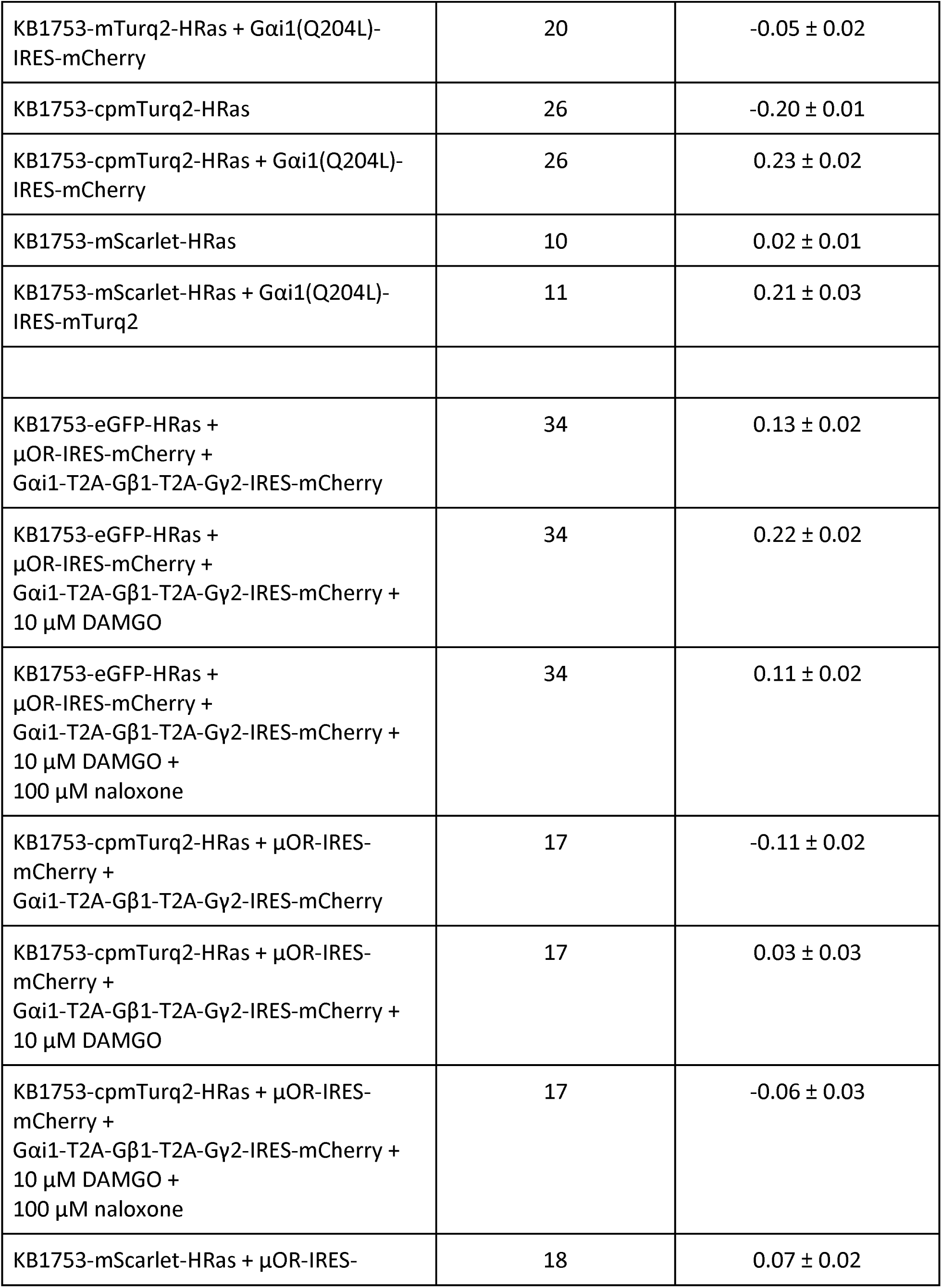
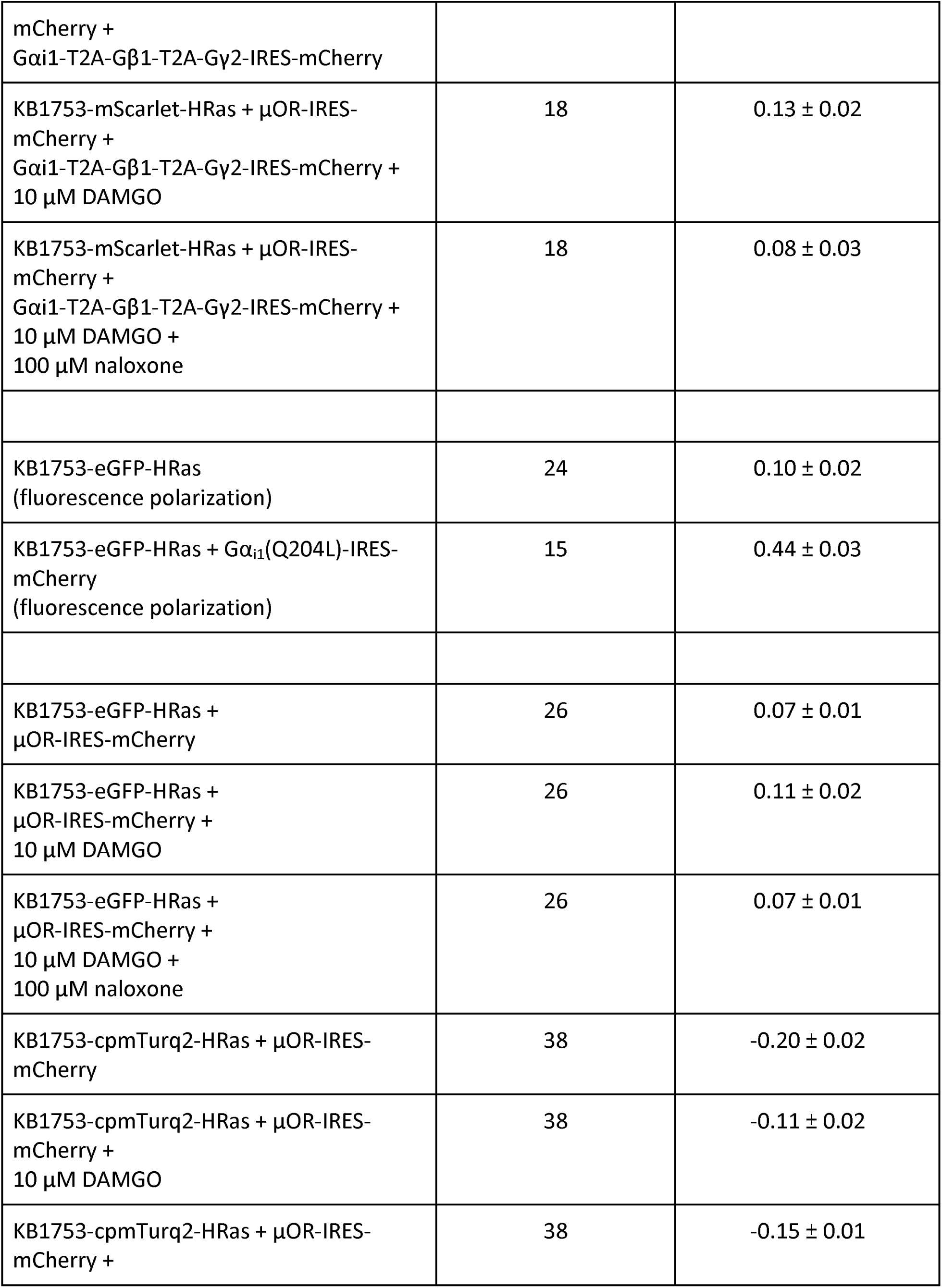
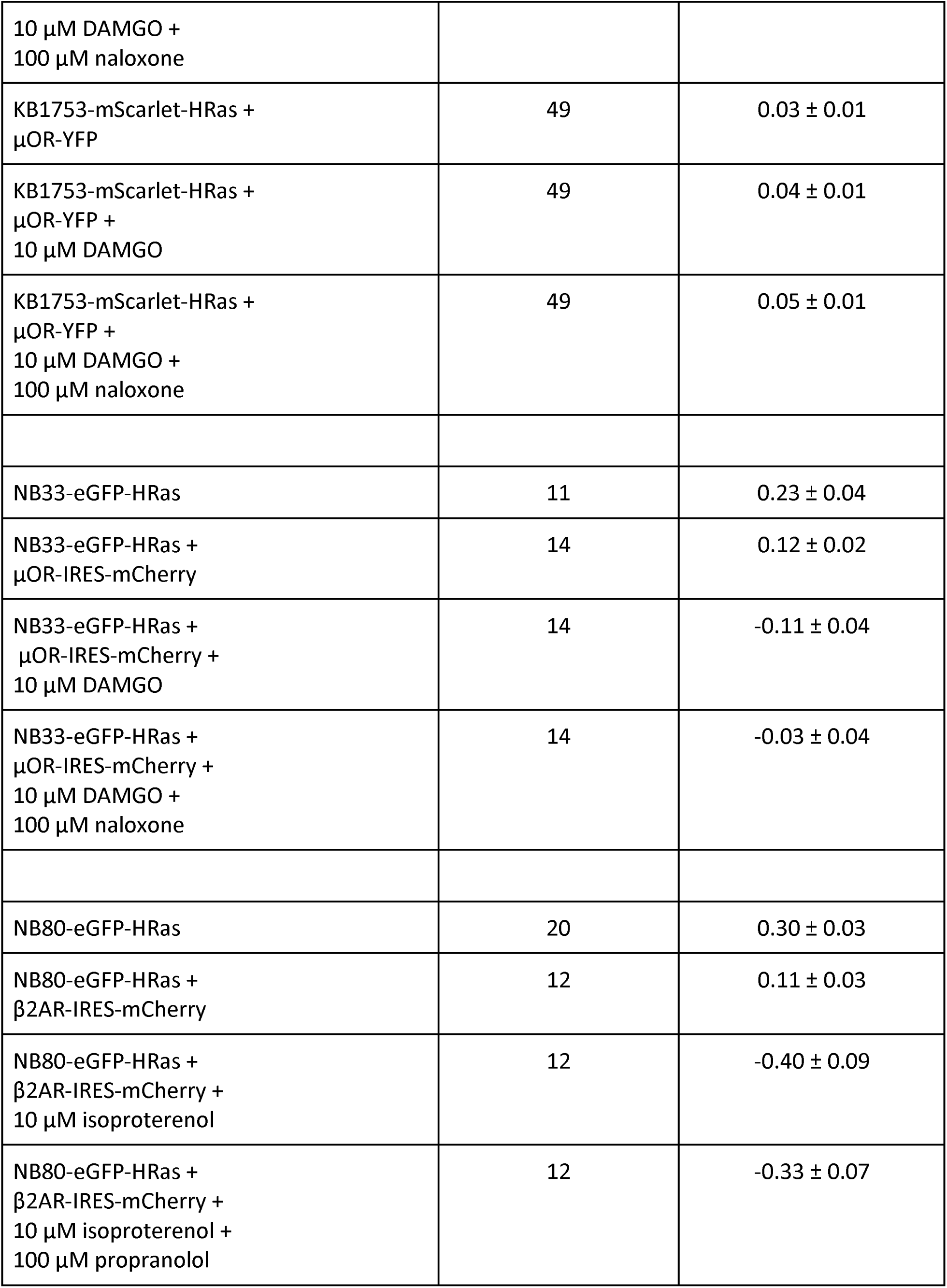
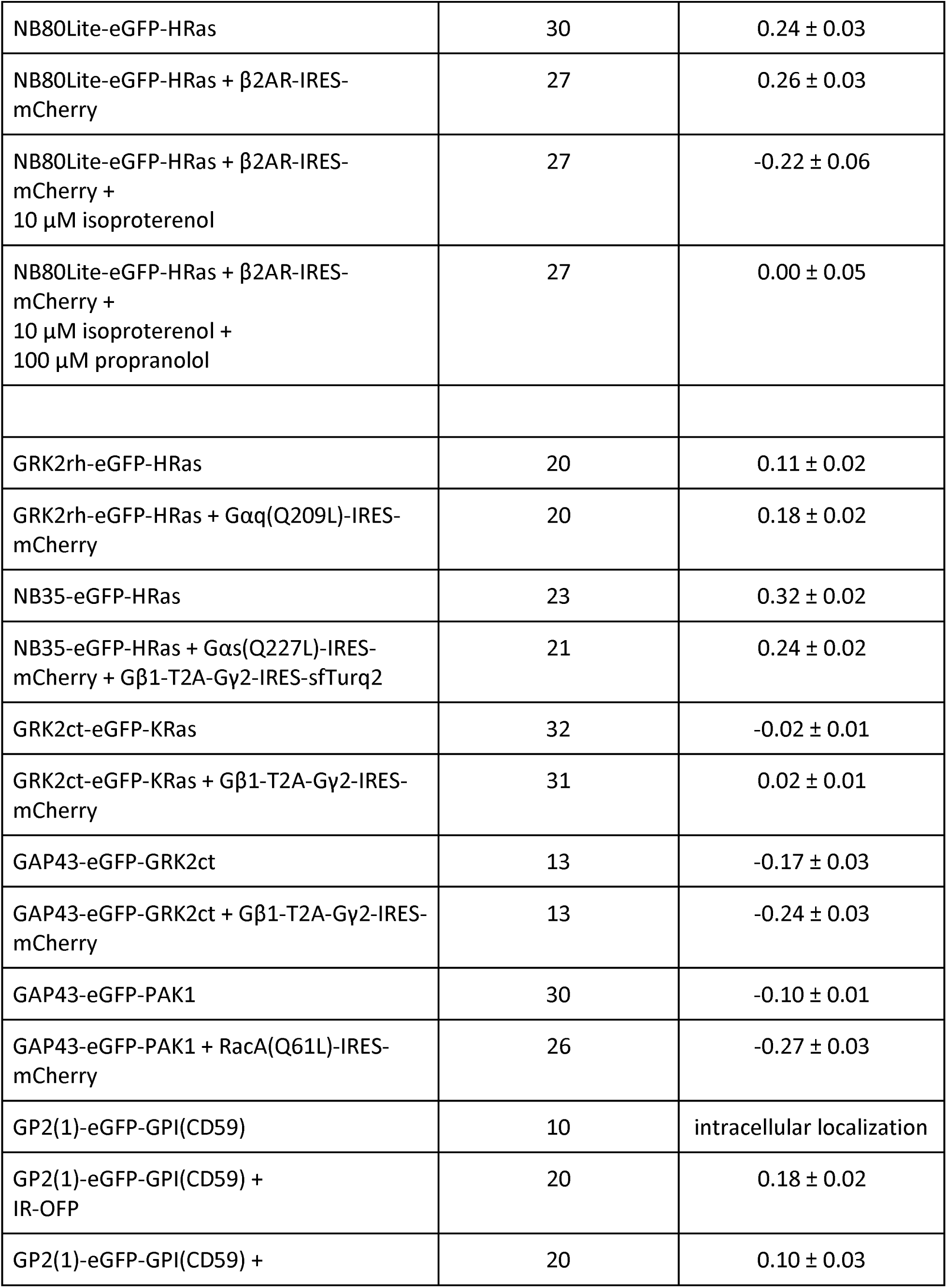
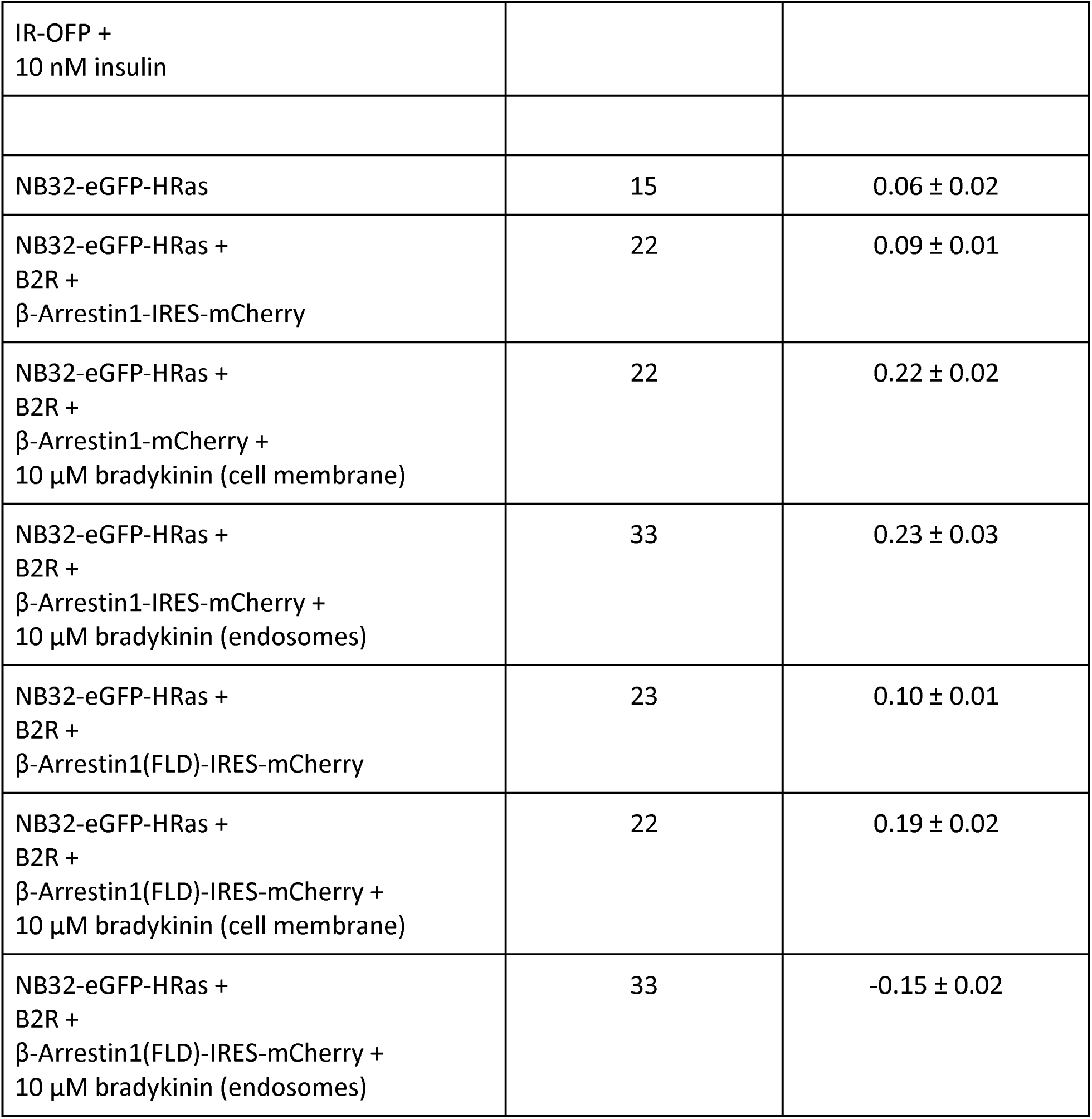

**Suppl. Fig. 1:**
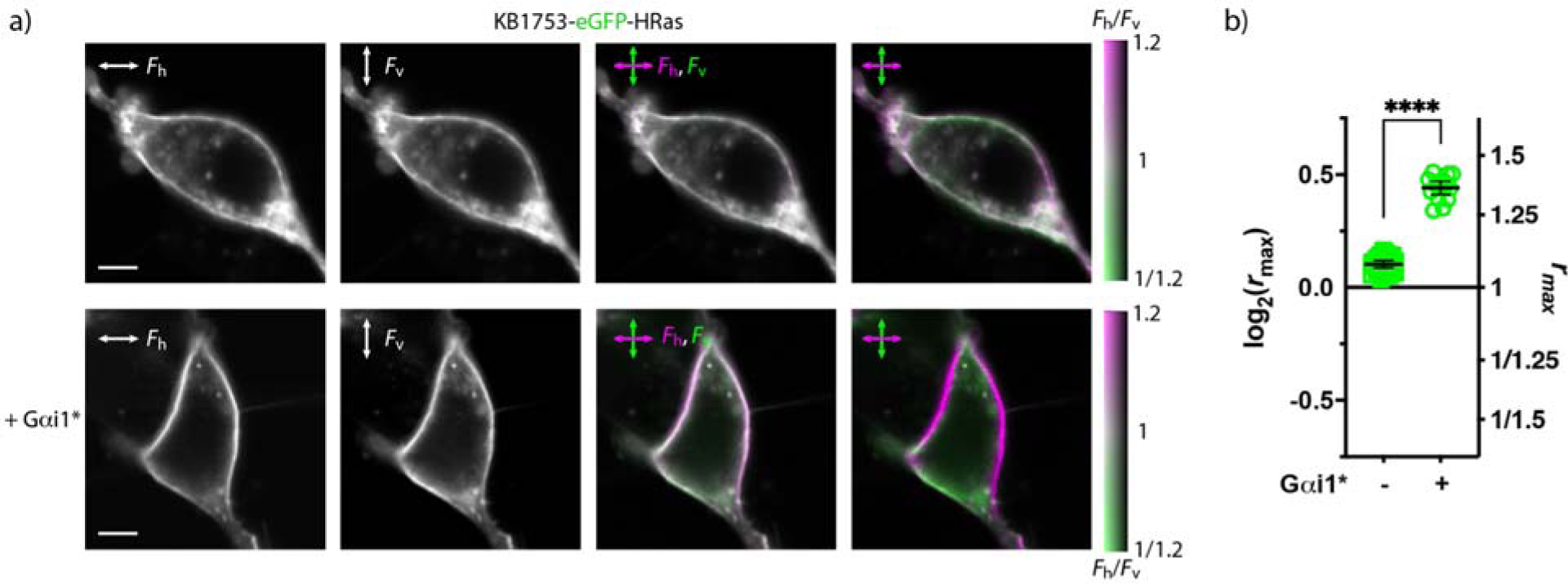
Fluorescence polarization imaging of HEK293 cells expressing KB1753-meGFP-HRas. **a)** A cell expressing KB1753-eGFP-HRas (top) and coexpressing KB1753-eGFP-HRas and Gαi1(Q204L)-IRES-mCherry (Gαi1*, bottom). From left to right: fluorescence polarized horizontally; fluorescence polarized vertically; the two preceding images colored magenta and green, respectively, and overlaid; same as in the preceding image, but the coloring adjusted to cover the shown range of fluorescence intensity ratios (F_h_/F_v_,), as indicated by the color bar. In cells expressing Gαi1*, the FLIP exhibits pronounced fluorescence polarization. Scale bars: 5 µm. **b)** Extent of fluorescence polarization in cells expressing KB1753-eGFP-HRas, in absence and presence of Gαi1*. Means and 95% confidence intervals are indicated. The probe exhibits significantly higher fluorescence polarization in presence of Gαi1*.

**Suppl. Fig. 2:**
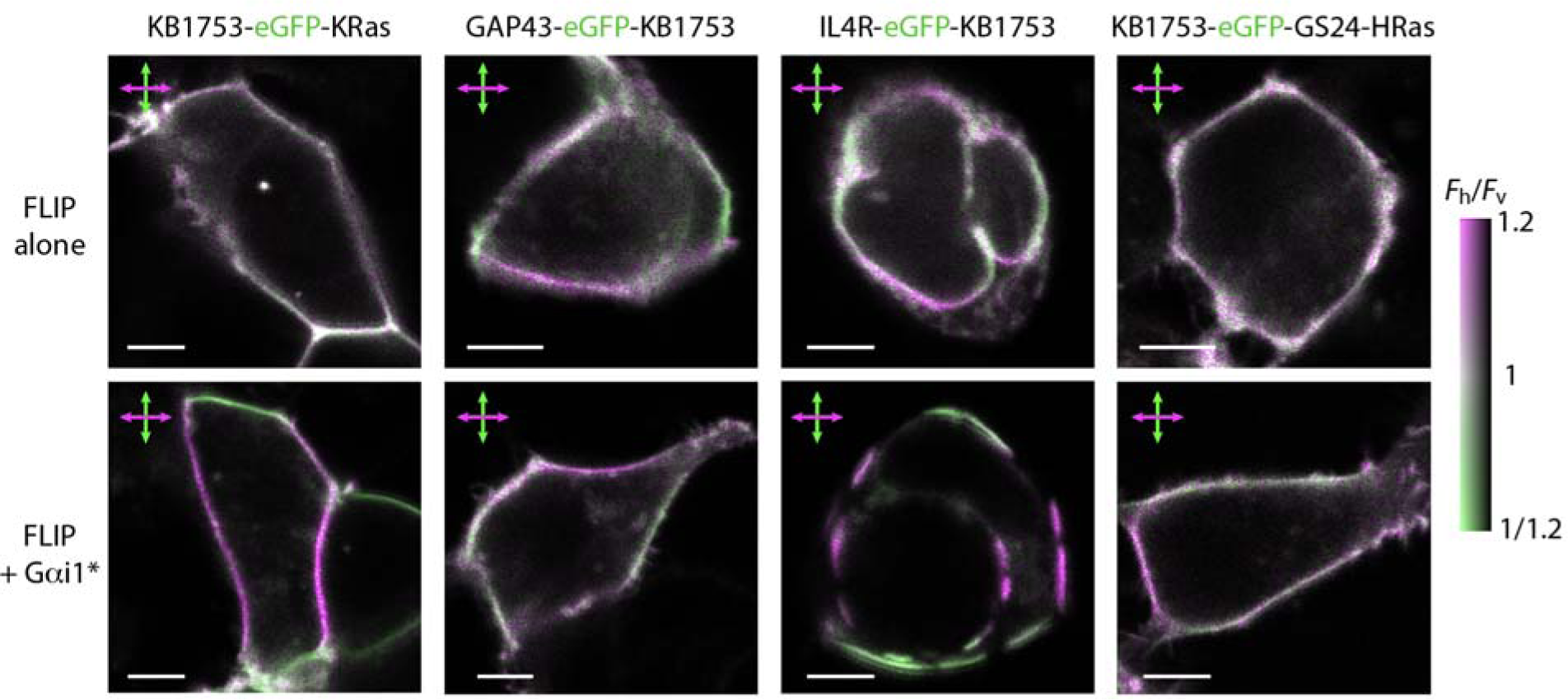
Effects of cell membrane anchors on LD of FLIPs containing the affinity binding peptide KB1753, in absence and in presence of a constitutively active mutant of Gαi1 (Gαi1*).

**Suppl. Fig. 3:**
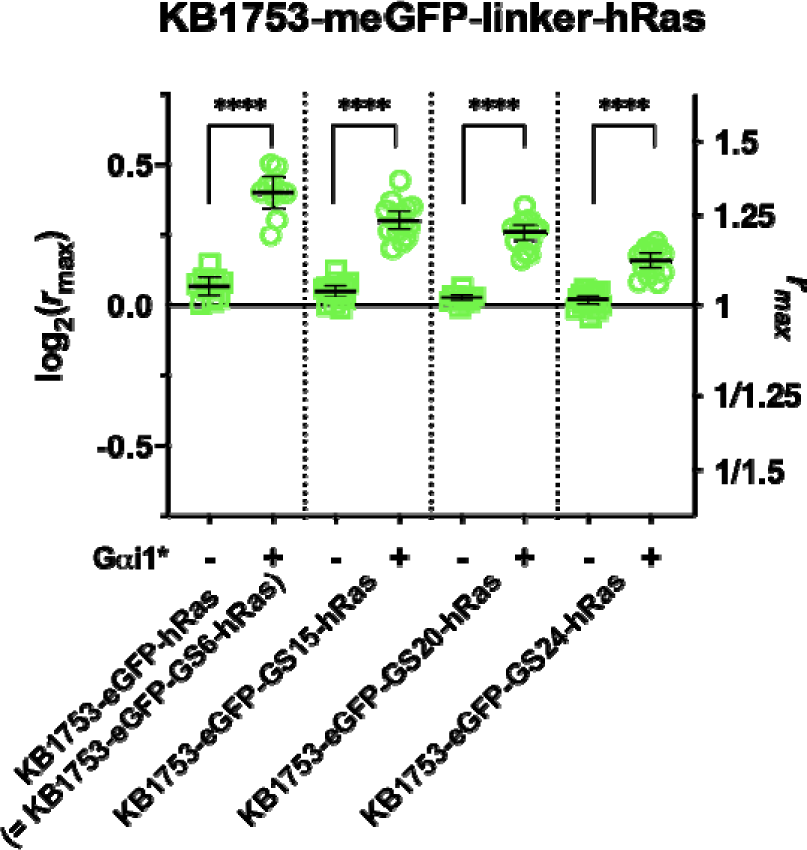
Effect of linker length in KB1753-meGFP-linker-HRas probes on their LD, in absence and in presence of a constitutively active mutant of Gαi1 (Gαi1*).

**Suppl. Fig. 4:**
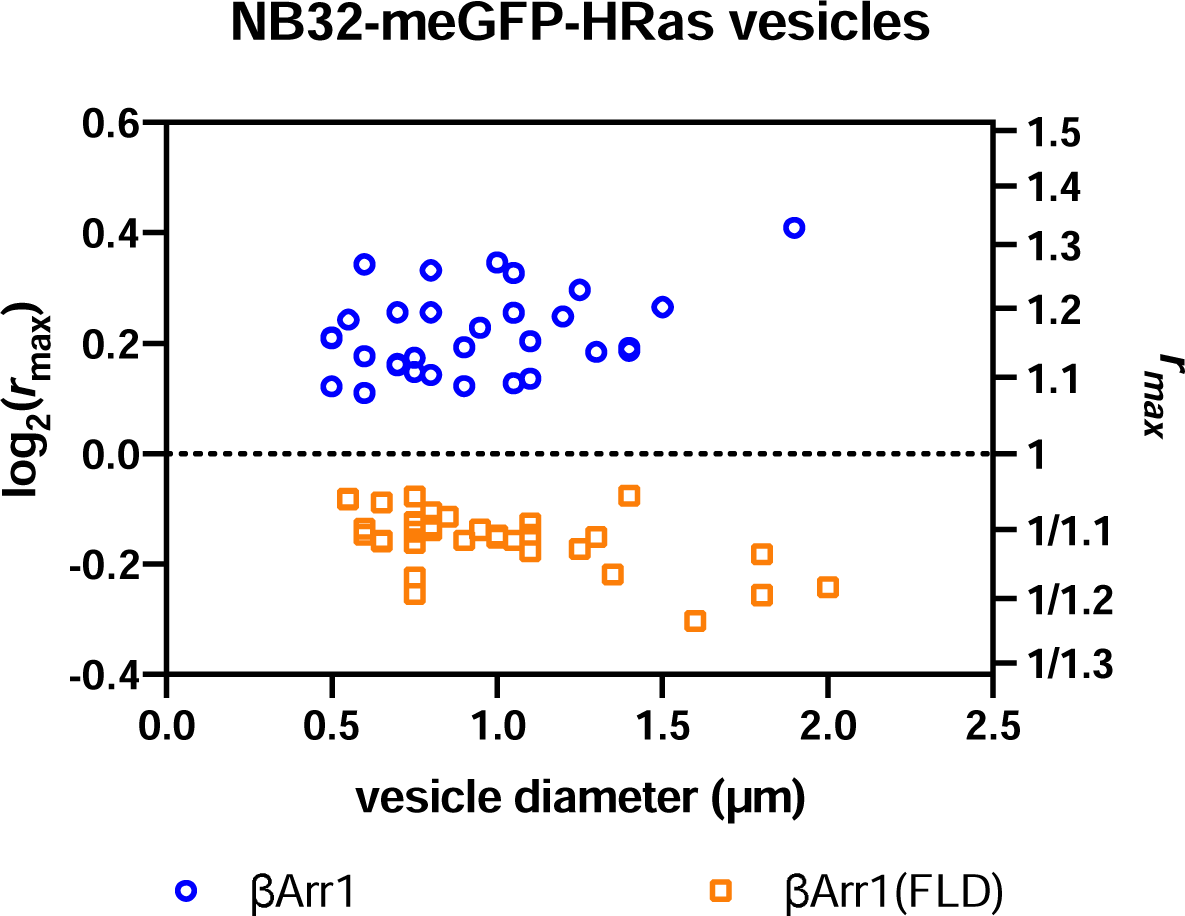
LD as a function of vesicle diameter in cells overexpressing B2R and βArr1-IRES-mCherry, or B2R and βArr1(FLD)-IRES-mCherry.

